# Nanopore sequencing of DNA concatemers reveals higher-order features of chromatin structure

**DOI:** 10.1101/833590

**Authors:** Netha Ulahannan, Matthew Pendleton, Aditya Deshpande, Stefan Schwenk, Julie M. Behr, Xiaoguang Dai, Carly Tyer, Priyesh Rughani, Sarah Kudman, Emily Adney, Huasong Tian, David Wilkes, Juan Miguel Mosquera, David Stoddart, Daniel J. Turner, Sissel Juul, Eoghan Harrington, Marcin Imielinski

**Affiliations:** Weill Cornell Medicine, New York, NY; New York Genome Center, New York, NY; Oxford Nanopore Technologies, New York, NY; Tri-institutional Ph.D. Program in Computational Biology and Medicine, New York, NY; Oxford Nanopore Technologies, Oxford, UK; Oxford Nanopore Technologies, San Francisco, CA

**Keywords:** Chromatin structure, Structural variation, Long read sequencing, *De novo* genome assembly, cancer genomics

## Abstract

Higher-order chromatin structure arises from the combinatorial physical interactions of many genomic loci. To investigate this aspect of genome architecture we developed Pore-C, which couples chromatin conformation capture with Oxford Nanopore Technologies (ONT) long reads to directly sequence multi-way chromatin contacts without amplification. In GM12878, we demonstrate that the pairwise interaction patterns implicit in Pore-C multi-way contacts are consistent with gold standard Hi-C pairwise contact maps at the compartment, TAD, and loop scales. In addition, Pore-C also detects higher-order chromatin structure at 18.5-fold higher efficiency and greater fidelity than SPRITE, a previously published higher-order chromatin profiling technology. We demonstrate Pore-C’s ability to detect and visualize multi-locus hubs associated with histone locus bodies and active / inactive nuclear compartments in GM12878. In the breast cancer cell line HCC1954, Pore-C contacts enable the reconstruction of complex and aneuploid rearranged alleles spanning multiple megabases and chromosomes. Finally, we apply Pore-C to generate a chromosome scale *de novo* assembly of the HG002 genome. Our results establish Pore-C as the most simple and scalable assay for the genome-wide assessment of combinatorial chromatin interactions, with additional applications for cancer rearrangement reconstruction and *de novo* genome assembly.

## Introduction

Mammalian chromatin is hierarchically organized into active and inactive compartments^1,2^, topologically associated domains (TADs)^3–5^, and loop domains^2,6,7^. A key challenge in epigenomics is to link multi-scale features of chromatin folding with the functional cellular programs that drive transcriptional regulation and cell identity^8,9^. An improved understanding of chromatin folding may also provide insight into the role of the epigenome in driving specific disease states. Previous work in constitutional genetics has linked the presence of inherited non-coding DNA variants, such as CTCF mutations^10^ or larger structural variants^11,12^, to human limb malformations through chromatin re-structuring^10^. In cancer, complex somatic genomic rearrangements can create neo-TADs^11,13^ or hijack enhancers to activate oncogenes^14–18^.

Our modern understanding of chromatin structure relies heavily on results obtained from chromosome conformation capture assays (3C, 4C, 5C and Hi-C) which utilize molecular readouts (e.g. PCR, sequencing) to map pairs of loci that spatially interact in the nucleus^1,19^. The general principles underlying these techniques can be modified to target specific nuclear proteins by immunoprecipitation (ChIA-PET^20^, HiChIP^21^) or specific loci using nucleic acid hybridization enrichment(ChiC^22^), and they can be adapted to study single-cell samples^23,24^. Most notably, Hi-C enables unbiased mapping of active (A) and inactive (B) chromatin compartments, TADs, cohesin loops, and enhancer-promoter interactions among the billions of interacting locus pairs in these genome-wide contact maps. In addition, Hi-C has been applied to generate reference-grade *de novo* assemblies of mammalian scale genomes^25^, identify structural variants in cancer cells^13^, and study DNA replication^26,27^.

Though analysis of Hi-C and its derivatives have revealed the unprecedented complexity of the 3D genome, the derived features are insufficient to explain many aspects of gene regulation^9,28–30^. The functional 3D state of chromatin, like other cellular machinery (e.g. protein signaling cascades), may not be fully described in terms of simple pairwise interactions. Rather, gene activation or silencing may require the interaction of three or more DNA loci, some of which may co-exist in dynamic and specialized nuclear structures that achieve cooperativity through liquid-liquid phase separation^31,32^. Detection of such higher-order chromatin complexes may be necessary to reveal fundamental links between genome structure and function.

Recent technologies enabling the study of three-way^33^ or higher-order chromatin interactions include chromosomal walks (c-walks)^34^, genome architecture mapping (GAM)^35^, split-pool recognition of interactions by tag extension (SPRITE)^36^, ChIA-drop^37^, Tri-C^38^, Tethered multiple 3C^39^, the concatemer ligation assay (COLA)^33^ and Multicontact 4C (MC-4C)^40^. Among these assays, only a subset (Tri-C, SPRITE, COLA, GAM) have been shown to generate genome-wide maps in mammals, and all methods but SPRITE suffer from very rare representation of (<1%) of contacts with order >3^36^. Though SPRITE contacts can reach a very high order (> 500), many may represent library artifacts arising from very large cross-linked fragments in the library preparation. As a result, they may serve as readouts of gross nuclear features rather than combinatorial chromatin states. In an ideal genome-wide higher-order chromatin assay, multi-way contacts would represent spatial adjacencies between sets of interacting DNA molecules (e.g. cooperative chromatin complexes arising during transcription factor binding).

Previous studies combining targeted chromatin conformation capture with long-read sequencing have revealed that many of the products of proximity ligation are in fact DNA “concatemers” of multiple interacting loci^33,34,37,40,41^. We surmised that chromatin conformation capture could be paired with nanopore sequencing to efficiently assay the combinatorial chromatin structure in human cells (**Fig. 1A**). We hypothesized that this approach would reveal higher-order 3D chromatin structure at both the scale of chromosomal compartments (i.e. multi-megabase) and chromatin loops (kilobase), identify novel 3D structures generated through complex cancer genomic rearrangements, and improve the scaffolding of long read, whole genome sequencing (WGS) derived assemblies.

**Fig. 1.**
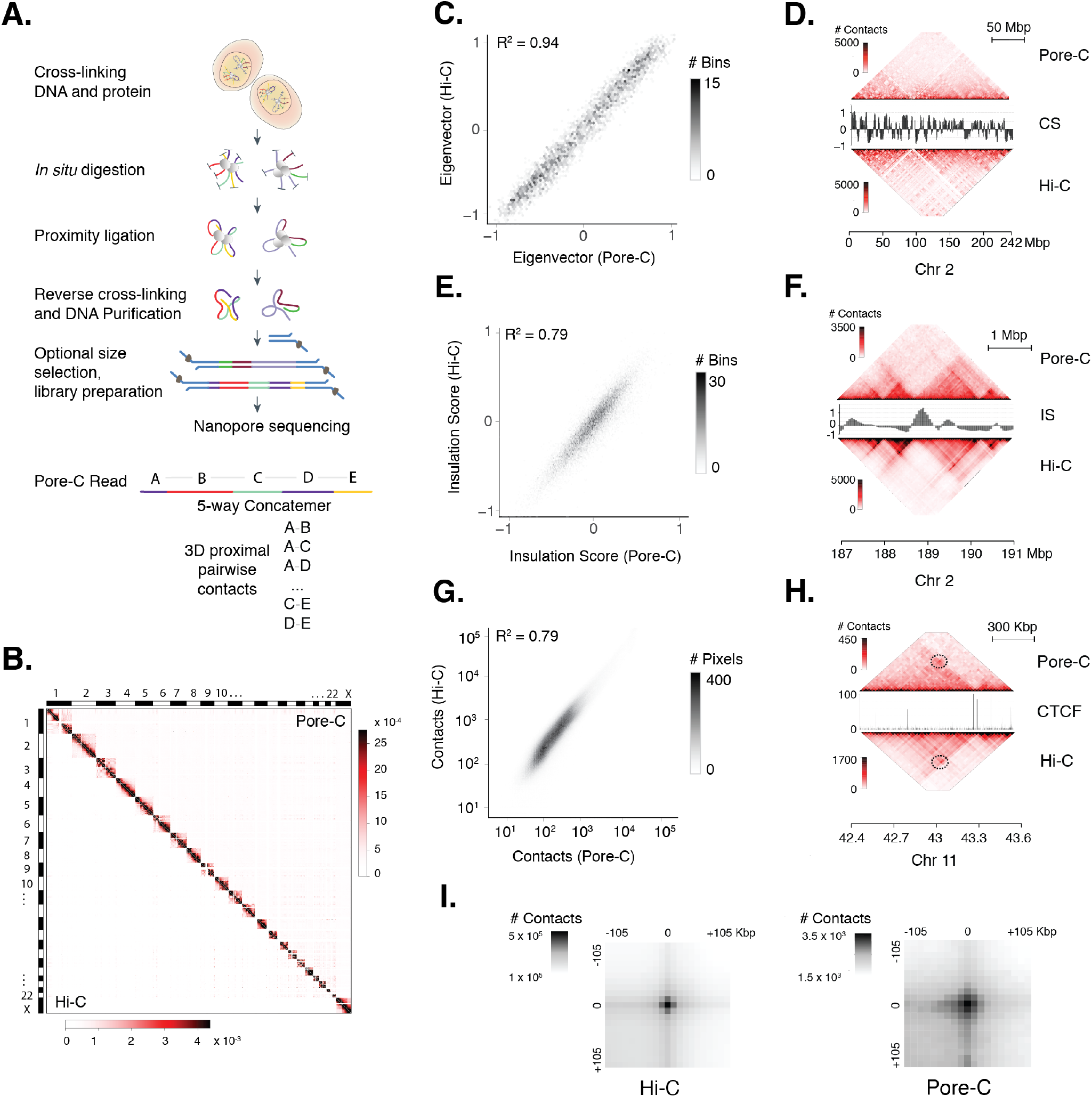
Pore-C concatemers reflect previously characterized features of chromatin architecture in GM12878. (A) Schematic of the Pore-C protocol: Proximally-ligated concatemers obtained by chromatin conformation capture are directly sequenced using Oxford Nanopore Technologies long-read sequencing platform. A greedy piece-wise alignment algorithm maps each concatemer sequence to a multi-way contact. (B) Comparison of Pore-C (upper triangle) 1 Mbp (virtual pairwise) and Hi-C contact maps^2^ (lower triangle) for chromosomes 1-22 and X (hg38) in the cell line GM12878. (C) Comparison of 500 Kbp compartment scores (CS) between Pore-C (1.38 billion virtual pairwise contacts) and Hi-C (~4 billion pairwise contacts) for GM12878^2^ (D) Comparison of Pore-C and Hi-C 500 Kbp contact maps for chromosome 2, alongside the CS demonstrating visual correspondence between Pore-C and Hi-C compartment patterns. (E) Comparison of 50 Kbp topologically-associated domain insulation scores (IS) between Pore-C and Hi-C. (F) Comparison of Pore-C and Hi-C 50 Kbp contact maps for a portion of chromosome 2, alongside the IS, demonstrating visual correspondence between Pore-C and Hi-C derived TAD structures. (G) Comparison of raw *cis* off-diagonal contact counts at 500 Kbp resolution between Pore-C and Hi-C. (H) An example of CTCF peaks coinciding with visually recognizable loop anchors (circle) in 25 Kbp Pore-C and Hi-C contact maps. (I) Results of aggregate peak analysis (APA) detecting similar enrichment of Pore-C and Hi-C pairwise data within 100 Kbp of loop anchors. In this map, each 10 Kbp x 10 Kbp pixel represents the total number of contacts detected across the entire loop set in a standard coordinate system centered around each loop anchor.

## Results

### Pore-C identifies higher-order contacts among DNA concatemers

We coupled nanopore sequencing with chromatin conformation capture to generate Pore-C libraries in an amplification-free manner through *in situ* digestion, and proximity ligation of cross-linked nuclei derived from the EBV-transformed B-lymphocyte cell lines GM12878 and HG002, as well as the breast cancer cell line HCC1954 (**Fig 1A**). For each sequenced Pore-C read, the Pore-C pipeline identifies a minimal subset of candidate alignments that maximally covers the entire Pore-C read. Each of these filtered alignments is then assigned to a restriction fragment based on the position of their midpoint (**Fig 1A, Fig S1A**). We refer to the set of filtered alignments associated with a given Pore-C read as a (multi-way) *contact*, and the number of fragments associated with a contact as its *order*.

To compare higher-order contacts with the pairwise contacts produced by Hi-C, we decompose each Pore-C read into all possible pairs, thus producing 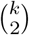 virtual pairwise contacts per read, where *k* is the order of the read (**Fig S1A**). Furthermore, we introduce the *pairwise contacts per Gbp* statistic as a measure of the contact density of the raw reads. In total we generated 597 Gbps of sequence data, detecting 145.2 million concatemers, 80.4 million of which had a contact order of 4 or more (**Table S1, Table S2**). These concatemers could be decomposed into 2.73 billion virtual pairwise contacts. We found that Pore-C libraries created using the NlaIII restriction enzyme had the highest contact density with an average of 6.5 million contacts per Gbp of sequenced data, closely matching the contact density of Hi-C datasets generated using 100 bp paired-end reads (**Fig S1D**) from the 4D nucleome project. We also observed that, although both the NlaIII and DpnII restriction enzymes were 4-base cutters, the fragment N50 of the NlaIII fragments in GRCh38 was roughly half the DpnII fragment N50 (**Fig S1B**) resulting in a lower contact density for the latter enzyme.

### Pore-C virtual pairwise contact maps are concordant with Hi-C

In order to compare Pore-C data to a previously-published “gold standard” GM12878 Hi-C dataset^2^, we constructed a contact map for GM12878 using NlaIII-derived virtual pairwise contacts (see above). Visual inspection of the Pore-C virtual contact map revealed previously identified features of GM12878 chromatin structure including A/B compartments (500 Kbp resolution, **Fig 1C**) associated with histone marks known to be predictive of active / inactive chromatin states (**Fig S2D**), topologically associated domains (TADs) (50 Kbp resolution, **Fig 1F**), and loops (10 Kbp resolution, **Fig 1H-I**), all of which were reflected in the corresponding Hi-C dataset. Additionally, we found a close correlation between Pore-C virtual pairwise contact maps and Hi-C data at the levels of eigenvector compartment scores (*R*^2^ = 0.94), TAD insulation scores (*R*^2^ = 0.79), and raw contact pixels (*R*^2^ = 0.79) (**Fig 1C, E and G**). To assess Pore-C run-to-run variability, the A/B compartment eigenvector score, TAD insulation score, and raw matrix correlations were computed for each of the individual NlaIII and DpnII runs (**Fig S2A-C**) and compared to the Hi-C dataset. We concluded that Pore-C DNA concatemers reflect known previously characterized pairwise features of chromatin architecture.

### High-order long-range contacts reflect combinatorial chromatin states

A considerable fraction of Pore-C concatemers represent multi-way (i.e. order > 2) chromatin contacts, with a median order of 7 (**Fig 2A**), and with 78.09%, 10.23%, and 1.04% of these reads harboring a contact order greater than 2, 10, and 20, respectively. In comparison, SPRITE clusters have a median order of 4 with 11.69%, 1.13%, and 0.4% of contacts with order greater than 2, 10, and 20, respectively. Additionally, we noted that SPRITE clusters were relatively depleted in multi-way contacts, and comprised a higher fraction of pairwise contacts (13.8%) and singletons (74.5%) compared to Pore-C (15.6% and 6.3%, respectively). This indicated a marked enrichment (OR = 18.5 for > 2 way; OR = 9.9 for > 10 way) of multi-way contacts in Pore-C as compared to SPRITE.

**Fig. 2.**
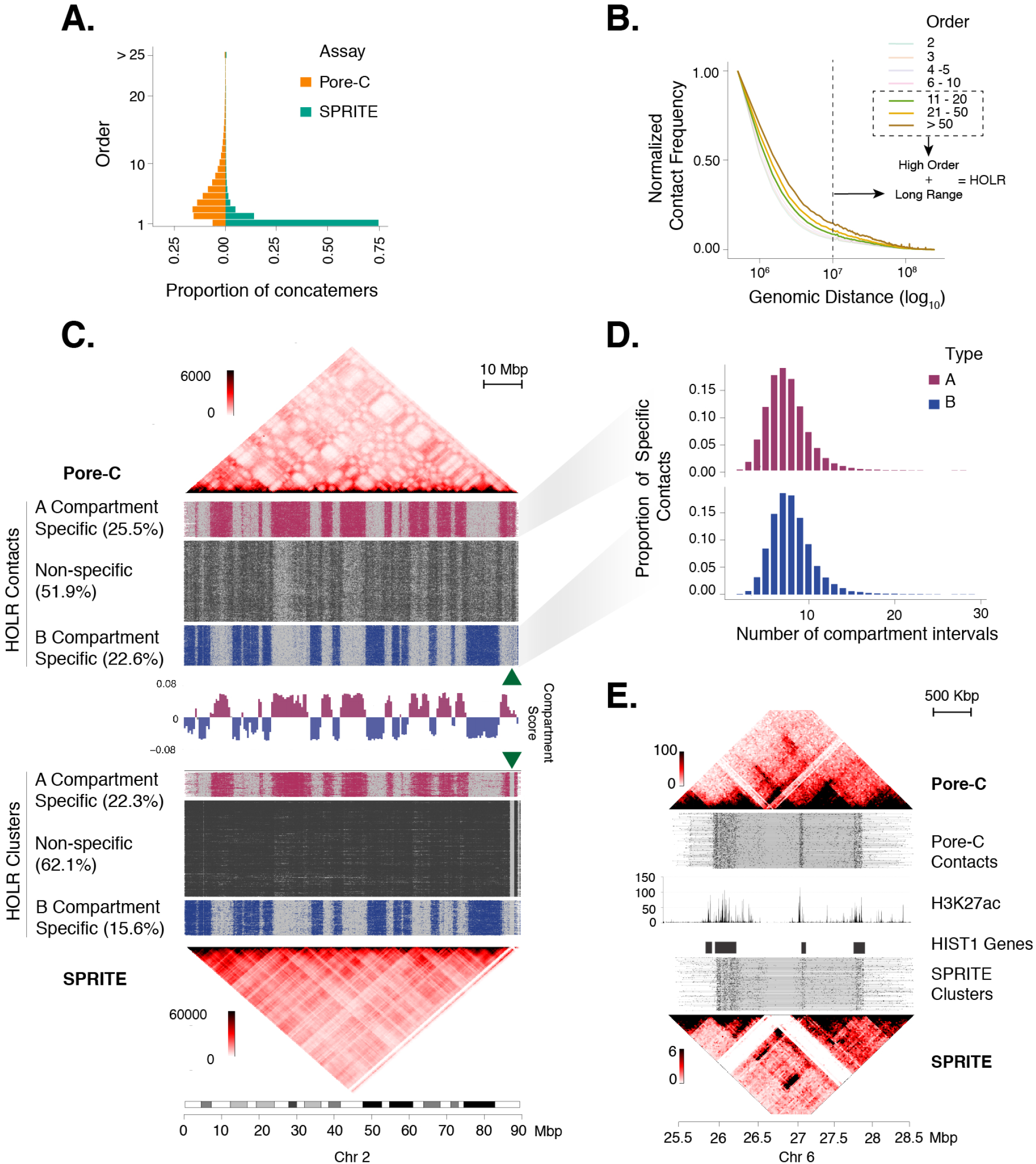
Pore-C concatemers reveal higher-order long-range contacts. (A) Comparison of the distribution of contact order between Pore-C concatemers and SPRITE clusters in GM12878. (B)Contact intensity as a function of linear genomic distance was plotted across all chromosomes for contacts of varying order. High-order concatemers (order > 10) showed a more gradual decay in contact intensity as a function of distance relative to lower orders. The plot depicts the thresholds for high-order (order > 10) long-range (distance > 10 Mbp) (HOLR) contacts used in the following panels. (C) HOLR contacts on chromosome 2p in GM12878 were called A-specific (>80% in A), B-specific (>80% in B), or non-specific according to their overlap with A and B compartment intervals. A and B compartment intervals were defined through segmentation of positive (A compartment) and negative (B compartment) values of the gold-standard Hi-C derived compartment score (middle track)^2^. HOLR contacts are plotted for these corresponding subgroups of Pore-C contacts (top half) and SPRITE cluster (bottom half) alignments. Percentages denote the frequency of A/B compartment-specific contacts among the set of HOLR contacts. These results show a higher fraction of A-specific and B-specific contacts in Pore-C, with additional combinatorial structure apparent in the non-specific Pore-C alignments that is less apparent in the corresponding non-specific SPRITE alignments. The pattern is also reflected in the virtual pairwise contact maps derived from Pore-C (top) and SPRITE (bottom) contacts. The green arrow indicates a peri-centromeric region that is absent in SPRITE but present in Pore-C due to its poor mappability with Illumina short reads. Giemsa banding pattern is shown below for scale and context. (D) Histograms of “compartment order” defined as the number of A or B compartments traversed by compartment specific Pore-C contacts. (E) Pore-C and SPRITE data are shown in the vicinity of a previously identified^36,42,43^ focal high-order histone locus body hub which spatially apposes three clusters of *HIST1* genes on human chromosome 6. Pore-C contacts (top track) and SPRITE clusters (bottom track) containing at least one fragment in at least one of the three *HIST1* gene clusters combinatorially connect peaks of active enhancer activity, as delineated by GM12878 H3K27ac ChIPseq (middle track). HOLR: High-order long-range

To explore the properties of multi-way contacts of increasing order, we divided this distribution into several groups based on order (2, 3, 4-5, 6-10, 11-20, 21-50, >50). We analyzed the genomic distance between pairs of monomers from decomposed concatemers, grouped by contact order (**Fig. 2B**). Previous studies have demonstrated high-order chromatin contacts connect distant genomic regions. This is reflected in a lower decay exponent *b* in the pairwise interaction probability *P* = *ad*^−*b*^ as a function of genomic distance *d* for multi-way vs. pairwise contacts^36,37^. To test this, we applied a generalized linear model to determine the effect of contact order on the decay exponent. We found that concatemers in the 11-20 (*P* = 0.00138, Wald test), 21-50 (*P* = 7.8 ×10^−8^), and >50 (*P* = 7.6 × 10^−14^) groups were associated with a significantly slower distance decay (**Fig. 2B**). We labeled these contacts (order > 10)as “high-order”. Upon inspection of the distance-contact plot we labeled contacts with distance greater than 10 Mbp as “long-range”.

To explore biological structures revealed by high-order and long-range (HOLR) contacts, we studied the combinatorial connectivity within and between A (active) and B (inactive) compartments revealed by both Pore-C and SPRITE (**Fig 2C**). Using A/B compartment definitions for GM12878 as defined by Rao *et al*^2^, we identified A (or B) compartment specific HOLR contacts as those with 80% alignment to the A (or B) compartment. In Pore-C concatemer libraries, 47.36% of HOLR contacts were compartment specific, with 19.78% and 27.58% demonstrating A- and B-specificity, respectively. In contrast, only 38.62% of SPRITE HOLR clusters demonstrated compartment specificity. We concluded that Pore-C HOLR contacts more accurately reflect direct molecular interactions relative to their SPRITE counterparts.

Plotting the alignment patterns of compartment-specific HOLR Pore-C contacts on chromosome 2p directly revealed the mutually exclusive A and B compartment anatomy (**Fig 2C**). Strikingly, compartment-specific Pore-C HOLR contacts were highly combinatorial, with the median contact intersecting 14 unique compartment intervals (range 2 - 29) with similar combinatorial complexity shown for A- and B– compartment-specific contacts (**Fig 2D**). Interestingly, visual inspection of the alignment patterns of non-compartment-specific HOLR Pore-C contacts revealed additional structures that most resembled B compartment alignments, but was distinct (**Fig 2C**). These results suggest that novel combinatorial compartment structures or inter-compartment dynamics may be revealed through the analysis of non-specific Pore-C HOLR contacts.

We found very subtle evidence of these additional structures among the 62.1 % of SPRITE HOLR clusters that were not compartment-specific; however, this signal was obscured by a substantial background of SPRITE clusters demonstrating uniform alignment along chromosome 2p, suggesting technical noise in the assay. Indeed, comparison of virtual pairwise contact maps across the region for Pore-C and SPRITE (across all concatemers / clusters) showed that SPRITE maps harbored a high burden of non-specific background (**Fig 2C**). In addition, SPRITE maps failed to resolve the compartment structure of a ~1 Mbp peri-centromeric region that is unmappable by short read sequencing (**Fig 2C**, green arrow). Pore-C HOLR alignments were able to firmly place this region into the active compartment of GM12878.

High-order chromatin structure is thought to be important to gene regulation and gene transcription, possibly through the induction of liquid-liquid phase separation enabling cooperative transcription factor kinetics^9,31,44^. Previously, SPRITE analyses provided evidence highof-order chromatin folding at a histone gene cluster on chromosome 6p, associated with the microscopically identifiable nuclear structure known as the histone locus body^36,42,43^. Compared to SPRITE, we observed clearer visualization of combinatorial connections between three clusters of histone genes (**Fig. 2E**) in Pore-C data. These foci of connectivity closely coincided with extended regions of H3K27ac (**Fig. 2E**), consistent with a chromatin configuration bringing multiple enhancers in 3D proximity. As with the compartment scale maps, Pore-C was also able to resolve structure in repetitive genomic regions that were unmappable and hence invisible to the (Illumia-based) SPRITE map (manifesting as white stripes in **Fig. 2E**).

### High-order chromatin contacts resolve the allelic structure of a complex cancer rearrangement

Structural DNA variants are common in cancer genomes, yielding complex loci harboring many copies of genomic intervals and rearrangement junctions in *cis* or *trans* phase^45,46^. To explore the phase of highly rearranged cancer loci, we profiled the breast cancer cell line HCC1954 with Pore-C, generating 97.7 Gb of sequence and 677.89 million pairwise contacts with NlaIII and 46.5 Gb of sequence and 211.0 pairwise contacts with DpnII. Analysis of publicly available short read Illumina whole genome sequence (WGS) data of the breast cancer cell line HCC1954 and its paired blood normal derived cell line HCC1954BL using SvAbA (local assembly based somatic junction caller)^47^, combined with our junction-balanced genome graph algorithm JaBbA^45^, identified a subgraph of amplified rearrangement junctions connecting three loci on chromosomes 9, 12, and 20. Amplified regions in these three loci comprise four discontiguous “islands” of copy number > 10, three of which span more than 300 Kbp.

Deconvolution of the junction-balanced genome graph into distinct alleles yielded multiple possible solutions that were consistent with junction and interval copy number inferred from short-read Illumina WGS. Visualization of the virtual pairwise contact map derived from the Pore-C library demonstrates long-range connectivity between these distinct amplified regions (**Fig 3A, top**). Integration of Pore-C high-order contacts with the possible alleles traversing this subgraph identified a candidate allele connecting these four intervals in *cis* as the most likely state (**Fig 3A, top**). This allele was supported by 1,213 Pore-C concatemers with order greater than 10, spanning two or more intervals on the allele, many of which were separated by long distances (>1 Mbp) on the inferred allele. To validate this complex allele, we performed three color metaphase FISH on HCC1954 and the diploid ANA51 cells, designing probes against three discontiguous genomic regions overlapping the reference genomic location of the inferred allele (**Fig 3B, bottom**). As predicted from our Pore-C data, these three probes were proximal and amplified in the HCC1954 cell line, and distinct and diploid in ANA51 (**Fig 3B, top**). These results confirm the presence of a complex and amplified derivative allele simultaneously connecting portions of chromosomes 9, 12, and 20 in HCC1954.

**Fig. 3.**
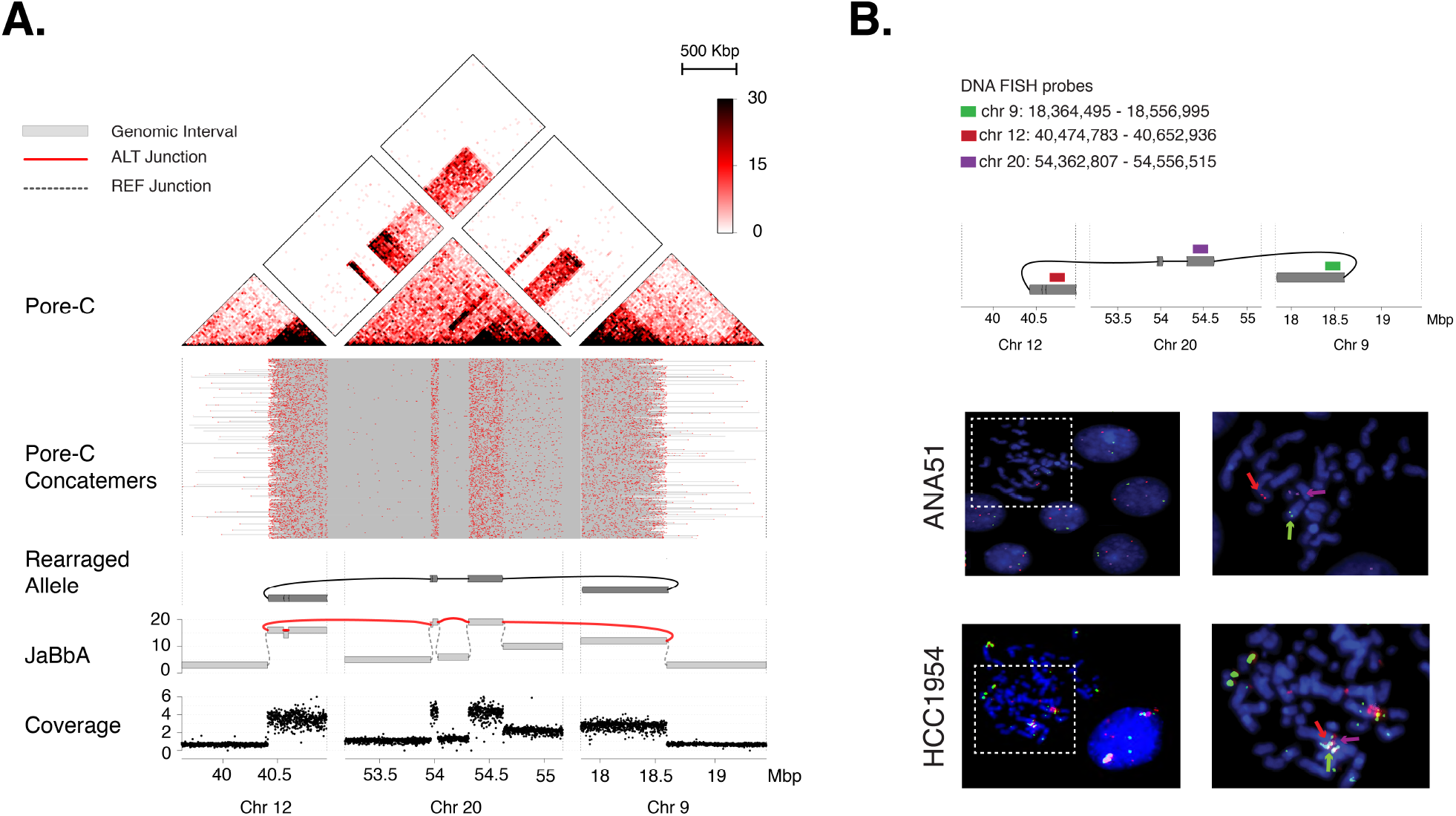
High-order chromatin contacts resolve the allelic phase of complex genomic rearrangements. (A) A complex, multi-chromosomal subgraph connecting chromosomes 9, 12 and 20 in the breast cancer cell line HCC1954 was identified using JaBbA, an algorithm for complex structural variant analysis (https://github.com/mskilab/JaBbA)^45^. Analysis of binned read-depth (bottom track) and junction calls yielded the genome graph (second track from bottom). At each edge, ALT represents an alternative junction that is depicted as a red line and REF represents a reference junction that is depicted as a gray dashed line. The gray bars each represent the genomic intervals of each connected node. Pore-C high-order contacts that mapped to the involved region (top track and heatmap) were used to nominate an amplified multi-junction derivative allele within this graph. The Pore-C alignments and associated contact maps (with each Pore-C read being highlighted in red and with gray lines representing the high-order connectivity between within each Pore-C contact) visually confirm this allele. (B) We designed a multi-color DNA FISH experiment (top) with red, green, and magenta probes each targeting one of three locations on this putative derivative allele. Co-localization of probes yielded a yellow signal in HCC1954 metaphase spreads (bottom), confirming the presence of an amplified multi-junction fusion of chromosomes 9, 12 and 20. We did not observe co-localization of these three probes in a diploid cell line (ANA51).

### Pore-C enables *de novo* assembly of chromosomal length scaffolds

In addition to the direct study of chromatin structure, chromatin conformation capture methods can be used to significantly improve the quality of *de novo* genome assemblies. Proximity information from Hi-C has been used to correct and merge contigs^50–52^ and even generate reference-grade mammalian genome sequences^25^. To assess the utility of Pore-C reads in scaffolding human genome assemblies, we analyzed the HG002 GIAB reference sample^53^. Whole-genome sequencing (WGS) libraries from HG002 high molecular weight genomic DNA were sequenced to roughly 40-fold average read depth with a read N50 of 22 Kbp using ONT’s promethION sequencing platform. We also generated three Pore-C libraries derived using the NlaIII, DpnII, and HindIII restriction enzymes. These were also run on the PromethION, and produced 218, 69, and 18 million virtual pairwise contacts, respectively.

The HG002 WGS dataset was processed using the `redbean` assembler which generated 2,374 contigs with an NG50 of 10.4 Mbp, with the longest contig spanning 82 Mbp (Fig.4A). We then applied `salsa2`^49^ to build scaffolds from these contigs. HindIII-derived Pore-C data yielded the longest scaffold (216 Mbp) with a nearly 10-fold increase in the sequence NG50 (98.6 Mbp) relative to the WGS data (**Fig. 4A**). Interestingly, although NlaIII-derived Pore-C libraries generated the highest number of high-order contacts (and hence pairwise contacts per Gbp), this data set only enabled a 3.2-fold improvement in the scaffold NG50 (33.2 Mbp) relative to the original assembly (**Fig. 4A**). DpnII demonstrated intermediate performance.

**Fig. 4.**
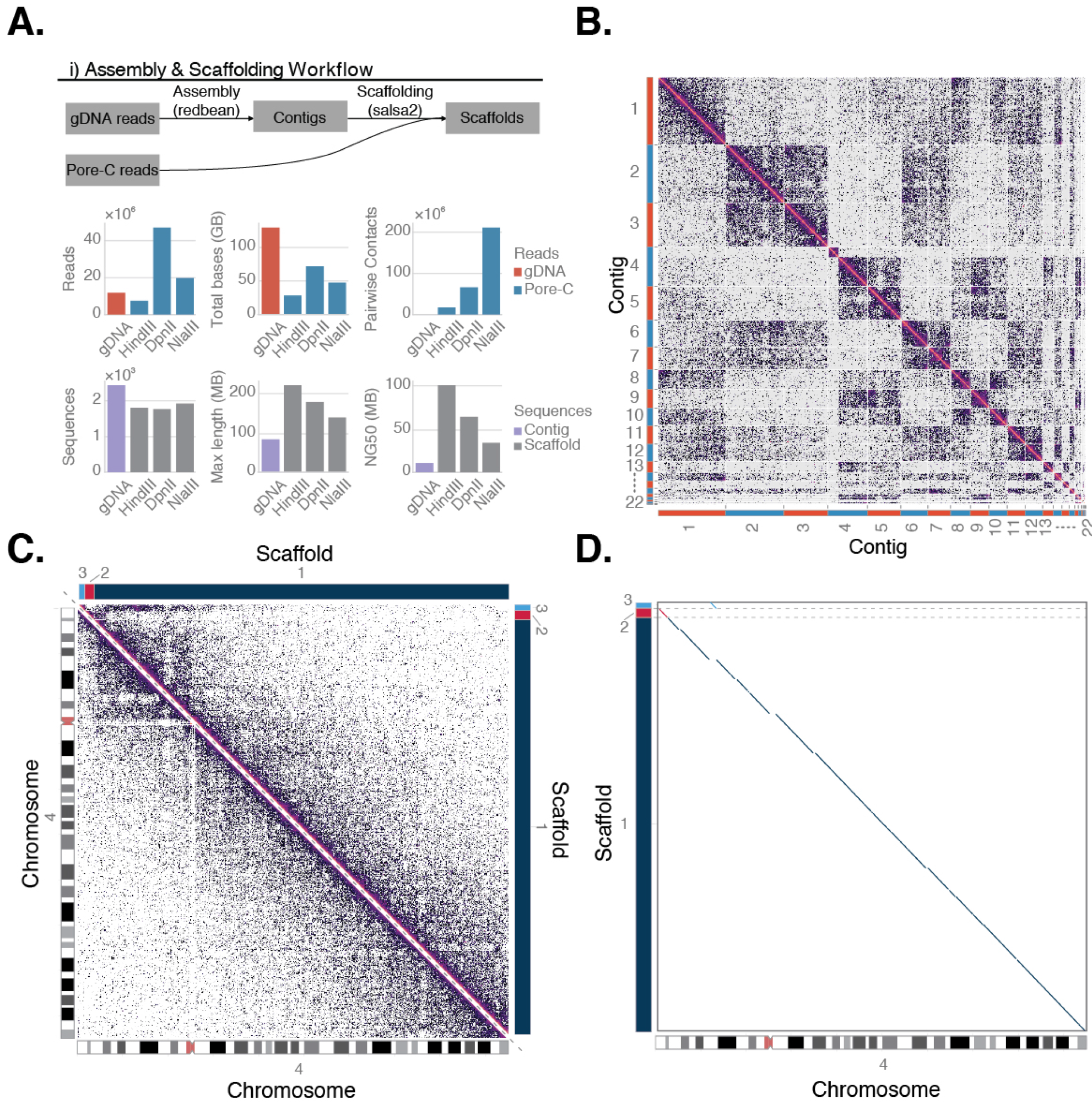
Pore-C concatemers guide assembly correction and scaffolding. (A) Assembly and scaffolding workflow was conducted in a 2-step process. ONT whole genome sequence data for HG002 was assembled using redbean^48^, then the resulting contigs were provided to `salsa2`^49^ along with pairwise Pore-C data generated from one of three different restriction enzymes to create a final scaffolded assembly. Despite having the fewest pairwise contacts the HindIII dataset created the most continuous scaffold. (B) Contact map showing the inputs to the scaffolding process for the HindIII dataset. Shown are the length-ordered contigs produced by redbean that map to chromosome 4. (C) Contact maps generated by mapping the HindIII Pore-C dataset against the scaffolds (above the diagonal) and the human reference (below the diagonal) for chromosome 4. The chromosome assembles into 3 scaffolds with 175 MB of 190MB in the largest scaffold, and although the centromere is not assembled, the two arms are properly placed in the scaffold relative to each other. (D) A dotplot of the scaffold aligned to GRCh38 shows that although there is a small misplaced scaffold, the assembly is largely correct.

Visualization of HindIII Pore-C data for sequences derived from chromosome 4 demonstrated a high-frequency of intercontig contacts (**Fig. 4B**). These were resolved following scaffolding, yielding a scaffold-scaffold contact map with the majority of interactions clustering at the diagonal (**Fig. 4C**). A dot plot revealed a high degree of synteny between HindIII Pore-C scaffolds and the hg38 chromosome 4 (**Fig. 4D**).

## Discussion

To our knowledge, Pore-C is the first assay to combine chromosome conformation capture with nanopore sequencing in an amplification-free manner. Our data also represent the only mammalian-scale demonstration of a genome-wide multi-way chromatin conformation capture assay that efficiently (i.e. in greater than 1 % of the data) detects contacts with order > 3. Though ChIA-Drop^37^ detects high-order contacts, it has been only been applied to *Drosophila melanogaster*, whose genome is an order of magnitude smaller than the human genome. SPRITE does not employ traditional chromatin conformation capture, but applies a ligation-free protocol. Relative to SPRITE, Pore-C shows a substantial (> 12 fold) enrichment of contacts with order > 3 (**Fig. 4D**). Other methods either have only been applied to the analysis of three-way interactions (Tri-C, GAM) and/or show rare representation of order > 3 contacts (COLA).

Through rigorous comparisons with Hi-C, we demonstrate high concordance between the probability of two sequences sharing a Pore-C concatemer and their Hi-C pairwise contact probability. As a result, Pore-C derived pairwise contact maps can be used to identify standard features of chromatin architecture, including A/B compartments, TADs, and loops. Based on our results, a single PromethION of a NlaIII Pore-C library can yield a contact map comprising >1 billion virtual pairwise contacts. These results are comparable to the yield from a ~2 billion read Illumina Hi-C library (based on the ~50% ligation frequency observed in Hi-C libraries). In this way, the Pore-C protocol represents a scalable alternative to a standard Hi-C experiment. Analytic caveats include the nonindependence of virtual pairwise contacts^33^ arising from the same concatemer, which may require correction (e.g. downweighting) when determining the effective resolution of a Pore-C contact map.

A key feature of Pore-C is its ability to detect high-order chromatin structure. Our analyses of contact order, A/B compartment connectivity, and histone enhancer hubs demonstrates similar or superior performance of Pore-C relative to SPRITE^36^. We demonstrate increased compartment specificity among Pore-C HOLR contacts relative to SPRITE, which when plotted along the genome allows direct visualization of the A or B compartment boundaries in GM12878. The complex high-order intra-compartment structure highlighted in this work will require further analytic exploration, including dissection of the combinatorial logic underlying the association of certain subsets of these multi-megabase reference genomic regions and their impact on gene regulation or cell identity. Intriguingly, we find additional structure among the ~50% of HOLR Pore-C contacts that do not show A or B compartment specificity. A key question is whether these additional patterns arise from distinct subpopulations of cells (e.g. cell cycle phases) or represent stochastic inter-compartmental crosstalk, like those that dynamically arise during gene regulation. Further algorithmic development and experimental work will be needed to explore these possibilities in Pore-C data, and nominate hubs of interactions (e.g. those arising from superenhancer-associated liquid liquid phase separations) genome-wide in a statistically calibrated manner.

The lower degree of compartment-specificity and overall combinatorial structure observed among HOLR SPRITE contacts relative to Pore-C may be due to technical features of SPRITE’s protocol. Since SPRITE employs ligation-free split-pool barcoding of variably-sized fragments of crosslinked chromatin, high-order SPRITE clusters may be effectively sampling very large spatial radius in the nucleus, traversing multiple compartments and/or chromosomes. Indeed, the combinatorial distance decay that we have shown higher-order Pore-C contacts is sharper than what has been previously shown for SPRITE or ChIA-drop^36,37^. By relying on the proximity ligation reaction, high-order Pore-C contacts may be less prone to connecting very spatially distant nuclear regions and thus may provide a more pure readout of spatial adjacency, and hence combinatorial chromatin complexes. Unlike droplet-microfluidic based approaches (ChIA-drop), Pore-C does not suffer from barcode collisions (i.e. from multiple library fragments occupying the same droplet and acquiring the same barcode) which can potentially inflate distance statistics, particularly for “high-order barcodes”.

Our results demonstrate the potential of high-order contacts to inform the reconstruction of complex rearranged cancer loci and *de novo* genome assembly. The data derived from high-order chromatin contacts in HCC1954 represent a sort of “ultra-long linked read”, in that each contact represents a set of (likely) contiguous sequences that are near each other in sequence. However, while 10X Chromium linked-reads are generated from fragments with average length of 50-80 Kbp, we show that high-order Pore-C contacts link multiple sequences across megabases of DNA. Furthermore, 10X Chromium linked-read libraries often suffer from “barcode collisions”, resulting from the fact 5-10 other fragments share a microfluidic droplet and the resulting barcode. These ambiguities require statistical criteria (i.e. multiple fragments spanning a structural variant breakpoint) and / or a minimally divergent reference genome (e.g. grouping read clouds according to mapping location) to resolve, which can impede complex structural variant detection or *de novo* assembly.

The vast majority of sequences contributing to proximally-ligated concatemers, and hence Pore-C contacts, arise in *cis* from a single contiguous molecule. As a result, we conjecture that Pore-C contacts will provide a less ambiguous source of long-range contiguity or phase information than 10X Chromium linked-reads. Though our assembly results demonstrate chromosomal length scaffolds from Pore-C using assembly tools that have been developed for Hi-C (`salsa2`), additional performance improvement will likely result from more specialized algorithms that fully leverage high-order Pore-C contacts to build longer scaffolds, as well as discern long-range phase blocks. Such algorithmic improvements may specifically impact the assembly and phasing of cancer genomes, where copy number alterations and rearrangements conspire to generate many copies of very similar alleles, or in high-ploidy domesticated plants such as wheat where broadly spanning phase information could link rare unique features throughout the genome, and significantly increase the genome assembly contiguity.

Unlike previous approaches combining proximity-based ligation with nanopore sequencing, the Pore-C protocol is amplification-free. As a result, the profiled sequences retain any DNA modifications (e.g. DNA methylation) that may be present on the input DNA molecule. Though we do not explore the analysis of base modifications on Pore-C reads, these signals could theoretically be used to probe the relationship between methylation states and combinatorial chromatin structures. Such an approach would serve as an alternative to Illumina-based bisulfite-sequencing based protocols for 3D methylation profiling (e.g. Hi-Culfite^54^, Methyl Hi-C^55^). Additional pipelines will be required to integrate methylation (and other base modifications detectable on nanopore reads) into the analysis of Pore-C contact data. We believe that the development of these and other algorithmic innovations will greatly expand the utility of this powerful and scalable assay for the study of high-order chromatin organization, complex cancer rearrangements, and *de novo* genome assembly.

## Methods

### Cell culture

Human B lymphocytes (GM12878) [Coriell Institute] and HG002 cells [Coriell Institute] were each maintained in suspension at a concentration greater than 200,000 cells/mL in RPMI-1640 [ATCC 30-2001] supplemented with 15% fetal bovine serum (FBS) and 1% penicillin/streptomycin. HCC1954 [ATCC CRL-2388] human breast cancer cells were maintained using a subculture ratio of 1:4 every three days in RPMI-1640 (ATCC 30-2001) supplemented with 10% FBS and 1% penicillin/streptomycin.

### Cross-linking

10 million cells were washed three times in chilled 1X phosphate buffered saline (PBS) in a 50 mL centrifuge tube, pelleted by centrifugation at 500xg for 5 min at 4°C between each wash. Cells were resuspended in 10 mL room temperature 1X PBS 1% formaladehyde [EMD Millipore cat no. 818708] by gently pipetting with a wide bore tip, then incubated at room temperature for 10 min. To quench the cross-linking reaction 527 μL of 2.5 M glycine was added to achieve a final concentration of 1% w/v or 125 mM in 10.5 mL. Cells were incubated for 5 min at room temperature followed by 10 min on ice. The cross-linked cells were pelleted by centrifugation at 500xg for 5 min at 4°C.

### Restriction enzyme digest

Each cell pellet was resuspended in a mixture of 50 μL of protease inhibitor cocktail [Sigma Aldrich cat no. P8340] in 500 μL of cold permeabilization buffer (10 mM Tris-HCl pH 8.0, 10 mM NaCl, 0.2% IGEPAL CA-630) and placed on ice for 15 min. Cells were centrifuged at 500xg for 10 min at 4°C after which the supernatant was aspirated and replaced with 200 μL of chilled 1.5X digestion reaction buffer [NEB] compatible with the restriction enzyme used. Cells were centrifuged again at 500xg for 10 min at 4°C, then aspirated and re-suspended in 300 μL of chilled 1.5X digestion reaction buffer. To denature the chromatin, 33.5 μL of 1% w/v SDS [Thermo Fisher Scientific cat no. 15553027] was added to each cell suspension and incubated for exactly 10 min at 65°C with gentle agitation then placed on ice immediately afterwards. To quench the SDS 37.5 μL of 10% v/v Triton X-100 [Sigma Aldrich cat no. 93443] was added for a final concentration of 1%, followed by incubation for 10 min on ice. Permeablized cells were then digested with a final concentration of 1 U/μL of either DpnII, NlaIII or HindIII [NEB] brought to volume with nuclease-free water to achieve a final 1X digestion reaction buffer in 450 μL. Cells were then mixed by gentle inversion. Cell suspensions were incubated in a thermomixer at 37°C for 18 hours with periodic < 1000 rpm rotation (< 30 sec every 15 min) to prevent condensation inside the lid.

### Proximity ligation and reverse cross-linking

DpnII and NlaIII restriction digests were heat inactivated at 65°C for 20 min with 300 rpm rotation. HindIII digests were chemically inactivated with final concentration of 0.1% w/v SDS at 65°C for 20 min with 300 rpm rotation, then quenched with a final concentration of 1% v/v Triton X-100 [Sigma Aldrich cat no. 93443]. Proximity ligation was set up at room temperature with the addition of the following reagents: 100 μL of 10X T4 DNA ligase buffer [NEB], 10 μL of 10 mg/mL BSA and 50 μL of T4 Ligase [NEB M0202L] in a total volume of 1000 μL with nuclease-free water. The ligation was cooled to 16°C and incubated for 6 hours with gentle rotation.

### Protein degradation and DNA purification

To reverse cross-link, samples were treated with 100 μL 20 mg/mL Proteinase K [Thermo Fisher Scientific cat no. 25530049], 100 μL 10% SDS [Thermo Fisher Scientific cat no. 15553027] and 500 μL 20% v/v Tween-20 [Sigma Aldrich cat no. P9416] in a total volume of 2000 μL with nuclease-free water. Samples were incubated in a thermomixer at 56°C for 18 hours with < 1000 rpm rotation (< 30 sec every 15 min) to prevent condensation inside the lid. In order to purify DNA, the sample was transferred to a 5 mL centrifuge tube, rinsing the original tube with a further 200 μL of nuclease-free H_2_O to collect any residual sample, bringing the total sample volume to 2.2 mL. DNA was then purified from the sample using a standard phenol chloroform extraction and ethanol precipitation.

### Nanopore sequencing

Purified DNA was SPRI size selected for fragments > 1.5 Kbp, then prepared for sequencing using either Oxford Nanopore Technologies SQK-LSK109, and sequenced on ONT’s MinION, GridION or PromethION platforms. In total, 27 sequencing runs were conducted generating a total of 449 Gbases of raw sequence from 3 different tissue sources including the human cell lines GM12878 and HG002 as well as the human breast cancer cell line HCC1954. The runs produced a total of 2.4 billion pairwise contacts. They were conducted using either DpnII, HindIII, or NlaIII for restriction enzymes in order to optimize the Pore-C methodology. The complete details for the runs are seen in (**Table S1**).

### Pore-C pipeline

We developed a reproducible analytic pipeline for deriving multi-way chromatin contacts from Pore-C concatemers (https://github.com/nanoporetech/pore-c/), and a workflow that can be found at (https://github.com/nanoporetech/Pore-C-Snakemake/) which uses the snakemake framework^56^. Briefly, the workflow involves alignment of Pore-C reads to a reference genome with Pore-C specific parameters. The resulting alignments are filtered to identify a minimal alignment set that covers each Pore-C read. Each of the resulting filtered alignments is then annotated with the set of reference genome restriction fragments with which it overlaps based on a *in-silico* restriction digest. From those fragment sets, each alignment is assigned to a single fragment based on the position of the alignment midpoint, which also serves to remove direct non-chimeric pairs that are caused by either incomplete digestion of the sample or ligation of cognate free ends. With the assignment of each alignment along a Pore-C read to a single restriction fragment, a multi-way contact is constituted, and this multi-way contact can then be decomposed into pairwise contacts. Decomposition to pairwise contacts involves enumerating the number of multi-way contacts containing a given pair of restriction fragments for every possible pair of restriction fragments in the genome. The fraction of alignments falling a given distance from the predicted in silico restriction sites in the GRCh38 reference genome were computed for DpnII, Nlalll and HindIII Pore-C datasets. The pairwise contacts are assigned to genome bins using conventional Hi-C workflows to generate a contact map, such as Hi-C explorer^57^ and cooler^58^. The workflow also includes an optional step to output the contact data in the bed format compatible with `salsa2`^49^ for genome assembly correction and scaffolding as well as the “medium text format” for conversion to the binary .hic format generated by the `juicer-tools` pre command (https://github.com/aidenlab/juicer/wiki/Download/) for use with the juicebox toolkit. Commands and parameters used for each step can be found in (table S2).

### Pore-C and Hi-C comparisons

Pore-C concatemers were decomposed into sets of virtual pairwise contacts in order to make correlations with existing Hi-C datasets. The resulting decomposed Pore-C contact matrix could be treated as a Hi-C pairwise contact matrix and explored using existing Hi-C analytic tools. The Pore-C aggregate dataset was composed of 11 NlaIII GM12878 Pore-C sequencing runs containing a total of 1.38 billion virtual pairwise contacts and the Hi-C dataset (4DNFIXP4QG5B)^2^ contained 4 billion contacts. Matrix balancing was performed on both the Pore-C and Hi-C contact matrices using the Knight-Ruiz iterative correction algorithm^59^ using `cooler balance`. A linear correlation between the Pore-C and Hi-C contact matrices measured three different similarity metrics i) raw contact matrices, ii) compartmental eigenvector scores, identified using `cooltools call-compartments` and iii) TAD insulation scores, calculated using the `cooltools diamond-arrowhead` tools. To further examine any variations between individual runs, the same three comparisons were made between each of the individual Pore-C runs, the complete Hi-C data set, and downsampled subsets of the Hi-C datasets. Downsampled Hi-C datasets were generated to match the contact count of each individual Pore-C run. Raw contact matrix, AB compartment and TAD correlations were performed between the down-sampled sets and the whole set to establish an upper bound for the correlation between any dataset of that size and the whole set. Publicly available ENCODE ChIP-seq peaks for H3K27ac, H3K4me3 and H3K4me1 (ENCSR447YYN) were used to visually confirm the correlation of Pore-C A/B compartmentalization with other existing markers of active / inactive chromatin structure. ENCODE CTCF ChiP-seq peaks (ENCSR000AKB) were utilized to identify loop anchors. The juicer suites APA tool was used to test for aggregate enrichment of Pore-C decomposed pairwise contacts within loop anchors identified in the Rao *et al*. GM12878 Hi-C dataset^2^.

### Analysis of high-order contacts

The change in pairwise contact frequencies as a function of genomic distance was determined by dividing concatemers into several groups based on order (2, 3, 4-5, 6-10, 11-20, 21-50, >50). The concatemers were then decomposed into pair-wise contacts and the normalized frequency of contacts as a function of genomic distance was plotted. This resulted into curves for each group. To statistically test the gradual decay of higher order contacts, a gamma-Poisson regression model was set up fitting raw frequency counts as: *count* ~ *log*(*total contacts*) + *group* + *log*(*distance*) with group comprising of 2-way contacts as the baseline. Wald-test from MASS R package^60^ was used to analyze significant deviations of different groups from baseline.

The high order long range (HOLR) contacts were defined if the maximum distance between the contacts for that concatemer, per chromosome, exceeded distance threshold of 10 Mbp (long range) and they belonged to group with order > 10 (high order).

For compartment-specificity analysis, each HOLR concatemer was assigned a label based on its overlap with A/B compartments defined using the compartment eigenvector scores derived from Hi-C dataset^2^ for the GM12878 cell line. The number of contacts falling in either compartment was tallied and proportions were calculated for each concatemer for each chromosome. Singleton contacts were filtered out. Focusing on multi-megabase level compartmentalization across the p-arm of chromosome 2, the performance of Pore-C to detect higher order interactions was assessed. For both Pore-C and SPRITE, a contact map using all Pore-C contacts or SPRITE clusters was created. Additionally, for each assay a read pile-up of Pore-C HOLR contacts or SPRITE HOLR clusters for A-specific, B-specific and unspecific HOLR contacts were plotted. The criteria of 80% or more contacts falling in a given compartment was used to determine compartment specificity. Lastly, compartment order for all HOLR A/B specific contacts were determined as the number of A or B compartments intersected by compartment specific Pore-C contacts.

For the analysis pertaining to *HIST1* gene cluster found on chromosome 6, previously described in^42^ and visualized using the SPRITE dataset^36^, both the SPRITE clusters and Pore-C contacts were filtered such that at least one fragment fell within one of these *HIST1* genes. The resulting read pileups for each technology were plotted along with the contact maps derived from each, as well as boxes to demarcate *HIST1* genomic locations.

We obtained GM12878 SPRITE data from (4DNFI-UOOYQC3)^36^. The tabular contact data was ingested using the R programming language in combination with a set of R/Bioconductor packages (including `GenomicRanges, rtracklayer, data.table`). All analyses of high order contacts were performed using Imielinski Lab R packages hosted at https://github.com/mskilab/ (`gUtils, bamUtils, gTRack, gGnome, GxG`) to represent, manipulate, analyze, and visualize genomic intervals, high-order contacts, and 1D / 2D genomic tracks.

### Validation of rearrangements in HCC1954 using Pore-C

Short read Illumina whole genome sequencing data for the breast cancer cell line HCC1954 and its matched normal HCC1954BL was obtained from the genomic data commons (GDC, https://gdc.cancer.gov/). DNA rearrangement junctions were detected using SvAbA^47^ and binned read depth was assessed using `fragCounter` (http://github.com/mskilab/fragCounter). Junction and read depth data were input into a junction-balanced genome graph inference algorithm JaBbA (https://github.com/mskilab/JaBbA)^45^. JaBbA integrates read depth and structural variation information for a given genome and produces a junction-balanced genome graph by solving a constrained optimization problem^45^. The junction-balanced graph allows querying of non-contiguous regions of genome that become connected through rearrangements through the R package `gGnome` (https://github.com/mskilab/gGnome). A sub-graph of amplified rearrangement junctions connecting three loci on chromosomes 9, 12, and 20 was identified. Using the sub-graph, all possible alleles that can give rise to the re-arrangement were combinatorially enumerated. Pore-C concatemer support was assessed for each derived allele. The allele with most concatemer support was selected and validated by FISH.

### Fluorescence *in situ* hybridization (FISH)

The fusion and amplification event detected between chromosomes 9, 12 and 20 were validated using three-color DNA fluorescence in situ hybridization (FISH). FISH probes were designed against loci of interest using the BAC clones(https://bacpacresources.org) RP11-341L12 (directly labeled green), RP11-476D10 (directly labeled red) and RP11-762A13 (directly labeled magenta). The HCC1954 and ANA51 cell lines were treated with Colcemid (0.1 g/ml) for 1 hour to obtain a metaphase preparation. Cells were then fixed in methanol/acetic acid (3:1). The fixed cells were then allowed to dry for 1 week on a microscope slide. Prepped slides were placed in a denaturation solution (70% formamide/2x SSC) for 15 minutes at 73°C proceeded by ethanol dehydration. Directly labeled probes were placed onto the slides and cells on the slide were then denatured for 15 minutes at 74°C proceeded by hybridization for 72 hours at 37°C. After hybridization, slides were washed in 2xSSC/0.3% NP-40, first at 73°C and then at 23°C. Metaphase cells were then stained with DAPI before visualization. A fusion-amplification event was measured as several green, red and magenta signals overlapping per nucleus or chromosome. Prior to use on HCC1954, all BAC clones were validated on metaphase spreads using the diploid ANA51 cell line. The nuclei and metaphase spreads were observed using a fluorescence microscope (Olympus BX51; Olympus Optical, Tokyo, Japan). Cytovision and Fiji software were used for imaging.

### Pore-C hybrid genome assembly

HG002 high-molecular weight (HMW) DNA was extracted from 20 million cells using the Puregene extraction kit [Qiagen cat. no 158667]. The sample was size-selected using the Circulomics Short Read Eliminator kit [Circulomics cat no. SS-100-101-01]. The HMW DNA was then sheared using a Megaruptor with a shearing speed of 28 and a library was prepared using a ONT library preparation kit [ONT cat no. SQK-LSK109]. The library was sequenced using the PromethION platform with nuclease flushes performed at 24 and 48 hours. Basecalling was performed using Guppy [version 3.2.2] using high accuracy mode. The sequencing run generated a total of 143.31 Gbp of sequence data with a read N50 of 22 Kbp. Reads shorter than 6 Kbp or with a mean read quality less than Q8 were removed, leaving 112.3 Gbp of raw sequence data. These reads were subjected to adapter trimming with PoreChop (https://github.com/rrwick/Porechop) and assembled with redbean^48^. This draft assembly was polished with 3 rounds of Racon^61^ followed by a round of Medaka polishing (https://github.com/nanoporetech/medaka). Pore-C libraries for each of the three restriction enzymes were generated from HG002 cells and sequenced on individual PromethION flow cells (**Table S2**). The virtual pairwise contacts for each enzyme were converted to a bed format compatible with the `salsa2`^49^ tool, which then converted the draft assembly to chromosome-scale scaffolds.

### Software Availability

Software used in this paper can be found in the following GitHub repositories:

https://github.com/nanoporetech/pore-c
https://github.com/mskilab/gGnome
https://github.com/mskilab/GxG

## ACKNOWLEDGEMENTS

We thank Jane Skok for helpful comments on the manuscript. Marcin Imielinski is supported by a Burroughs Wellcome Fund Career Award for Medical Scientists, Doris Duke Clinical Foundation Clinical Scientist Development Award, Starr Cancer Consortium Award, Melanoma Research Alliance Team Science Award, and National Institutes of Health U24-CA15020.

## AUTHOR CONTRIBUTIONS

These contributions follow the Contributor Roles Taxonomy guidelines: https://casrai.org/credit/. Conceptualization: N.U., M.I.; Data curation: N.U, M.P.,E.H., M.I.; Formal analysis: N.U., M.P., A.D., E.H., M.I.; Funding acquisition: S.J.,D.T.,M.I.;Investigation:N.U.,M.P.,A.D.,S.S.,X.D.,E.H.,S.J.,M.I.Methodology: N.U., M.P, A.D., S.S, D.S., C.T., S.J., E.H., M.I.; Project administration: E.H., M.I.; Resources: S.J, M.I.; Software: N.U., M.P., E.H., A.D., M.I. Supervision: S.J., M.I.; Validation: N.U., M.P., A.D, X.D., S.K., E.A., D.W., J.M.M, E.H., S.J., M.I. Visualization: N.U., M.P., A.D., M.I. Writing – original draft: N.U., M.P., M.I. Writing – review & editing: all authors.

## Supplementary Material

### Glossary

*Virtual Digest*: A genome sequence informatically fragmented at every restriction site for an enzyme and represented as a set of non-overlapping intervals along the genome.

*Virtual Contact Map*: A contact map generated from multi-way contact data by counting the number of multi-way contacts supporting the co-observation of each member of every pair of contacts.

*Restriction Fragment*: A single interval produced by a restriction digest.

*Pore-C read*: An ONT read generated by sequencing the DNA from a Pore-C library.

*Pore-C read alignment*: Alignment of Pore-C reads against a genome sequence. Specialized alignment parameters must be be used due to the scrambled nature of the concatemer.

*Fragment Assignment*: The process by which intervals along a Pore-C read are mapped back to restriction fragments produced by the virtual digest. Can be thought of as a bioinformatic method of decoding what happened during the digestion and ligation steps of the *pore-C protocol*. Spurious alignments must be filtered out during this step.

*Pore-C singleton*: A Pore-C read where only a single restriction fragment can be assigned. This contains no proximity information and is thus filtered out.

*Pore-C multimer*: A Pore-C read where multiple restriction fragments can be assigned.

*Pore-C pair*: A pair of restriction fragments present in the same Pore-C multimer. The standard Pore-C analysis uses all possible pairs of restriction fragments.

*Direct pair*: a pair of restriction fragments that are adjacent on the read and is roughly equivalent to what a chimeric Hi-C read detects.

*Indirect pair*: a pair of restriction fragments that are not adjacent on the read (i.e. those with other restriction fragments between them).

*Concatemer*: A set of restriction fragments ligated together into a single molecule by the Pore-C protocol. Restriction fragments that tend to be close in 3D space will tend to be part of the same concatemer.

*Monomer*: A single restriction fragment in the context of a concatemer.

*Contact order*: The number of participating restriction fragments in a single multi-way contact. A single Pore-C run will generate a distribution of contact orders depending on restriction enzyme motif frequency and the read length distribution of the run itself. *Chimeric Junction*: A junction in a concatemer generated by the ligation of two restriction fragment ends that are non-adjacent in the genome. These junctions are used to infer 3D proximity.

*Non-Chimeric Junction*: A junction in a concatemer generated by the ligation of two restriction fragment ends that are adjacent in the genome. This can be the result of incomplete digestion, the re-ligation of cognate ends of a digestion event or possibly a SNP at the digestion site.

*high order long range (HOLR)*: contacts defined as a multi-way contact containing at least 10 monomers in *cis* whose maximum span is greater than 10 megabases.

### Pore-C pipeline

Alignment of raw sequencing data was conducted with bwa using parameters outlined in (**Table S2**). The resulting alignments were then filtered to a subset of alignments that maximally cover the read. Filtering was done using a directed acyclic graph (DAG) traversal formulation of the alignment coverage problem. For each read, a DAG is constituted containing one node for each alignment and two additional nodes which are named “START” and “END”. Graph edges are added from START to each of the alignments such that the value of the edge is equal to the bwa bwasw gap penalty for a gap whose size is equal to the distance from the start of the read to the start of the alignment minus the alignment score. Additional edges are added from each alignment to the END node whose values are equal to the bwa bwasw gap penalty of a gap equal to the distance from the end of the alignment to the end of the read. Edges are then added between every pair of alignments A,B such that the edge source is the alignment whose endpoint is closer to the beginning of the read and the sink is the alignment whose endpoint is closer to the end of the read. The edge value is determined based on four relational classes: non-overlapping alignments, dovetailed alignments, contained/containing alignments, co-terminal alignments. First assume that the end of alignment A is closer to the beginning of the read than B. If alignments A and B do not overlap, then the edge value is equal to the gap penalty of the distance from the end of A to the start of B minus the alignment score of B. If A and B do overlap, then the edge value is equal to the gap penalty of an interval of size start of B to the end of A. If alignment A is contained within alignment B, then the edge score from A to B is computed similar to if A and B simply overlap. If a pair of alignments has the same endpoint, then the edge is drawn from the alignment whose start is closer to the start of the read to the second alignment in the pair and the value is equal to the alignment score of the shorter alignment plus a gap penalty the length of the second alignment. The graph is then traversed from the START node to the END node using bellman-ford algorithm to find the collection and order of alignments that has the optimal aggregate alignment score, and therefore the greatest non-redundant coverage of the sequencing read (**Fig S1A**). In addition to removing alignments that are not part of the optimal alignment tilepath, low quality alignments are removed, and additional filtering is done to remove singleton alignments, where a read has only a single alignment and is therefore unable to constitute even a single pairwise contact. For the purpose of creating a balanced contact matrix, we found it necessary to use a balancing blacklist https://rdrr.io/bioc/HiCcompare/man/hg38_blacklist.html. While Pore-C read monomers can confidently align to regions that are generally inaccessible to Hi-C, these regions are often represented by model sequences that differ, by design, in copy number from true biological samples. This results in artificial pileups of alignments that prevent proper matrix balancing and downstream analysis.

**Fig. S1.**
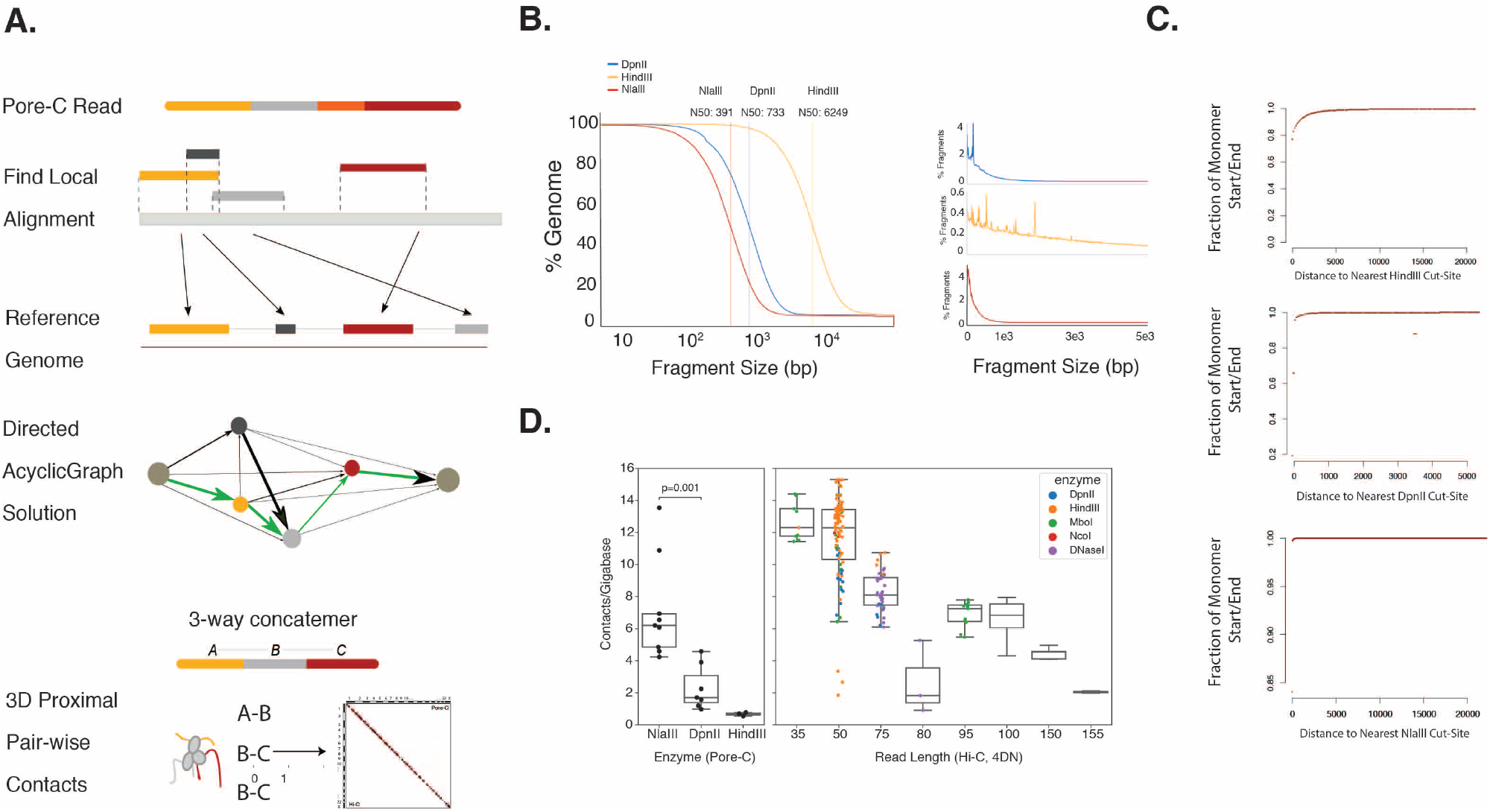
Optimization of Pore-C Methodology. (A) An overview of the directed acyclic graph filtering algorithm. A read is aligned to a reference genome. The DAG is constituted first with nodes representing the read start and end, and then with edges reflecting the combination of the alignment score as well as a gap or overlap penalty between the two involved reads or a given read and either the start or end of the read. Graph traversal from the start to end nodes describes a combination of alignments that maximally cover the read. (B) Comparison of the fragment length distributions generated by the three different restriction enzymes in the human reference GRCh38. (C) The fraction of reads falling a given distance from predicted restriction sites were calculated for each restriction enzyme. Although alignments generated with our workflow are not filtered based on whether they terminate close to a restriction site, alignment ends tend to fall very close to predicted sites. (D) By calculating the number of pairwise contacts generated per Gbp sequenced, head to head comparisons can be made with Hi-C, showing that Pore-C using NlaIII can output roughly as many contacts per gigabase sequenced as 100 bp paired-end Illumina Hi-C reads.

### Data quality assessment

Practically speaking, read N50 can be used as a simple indicator of a ligation efficiency quality metric for a given library. As a final metric we also calculate “pairwise contacts per Gbp sequenced”, which enables users to determine the cost efficiency compared to Hi-C. When compared directly to Hi-C runs from 4DN, NlaIII runs generate a similar contact density to 100 bp paired-end Hi-C runs and approach the contact density of 35 bp paired-end Hi-C runs (**Fig S1B**). Finally, because our published pipeline does not consider the predicted locations of restriction sites in the reference genome as prior data for assignment of alignments to restriction fragments or for filtering, the robustness of our filtering protocol is affirmed by the fact that most of the ends of alignments are located very close to predicted restriction sites (**Fig S1C**).

**Table S1.**
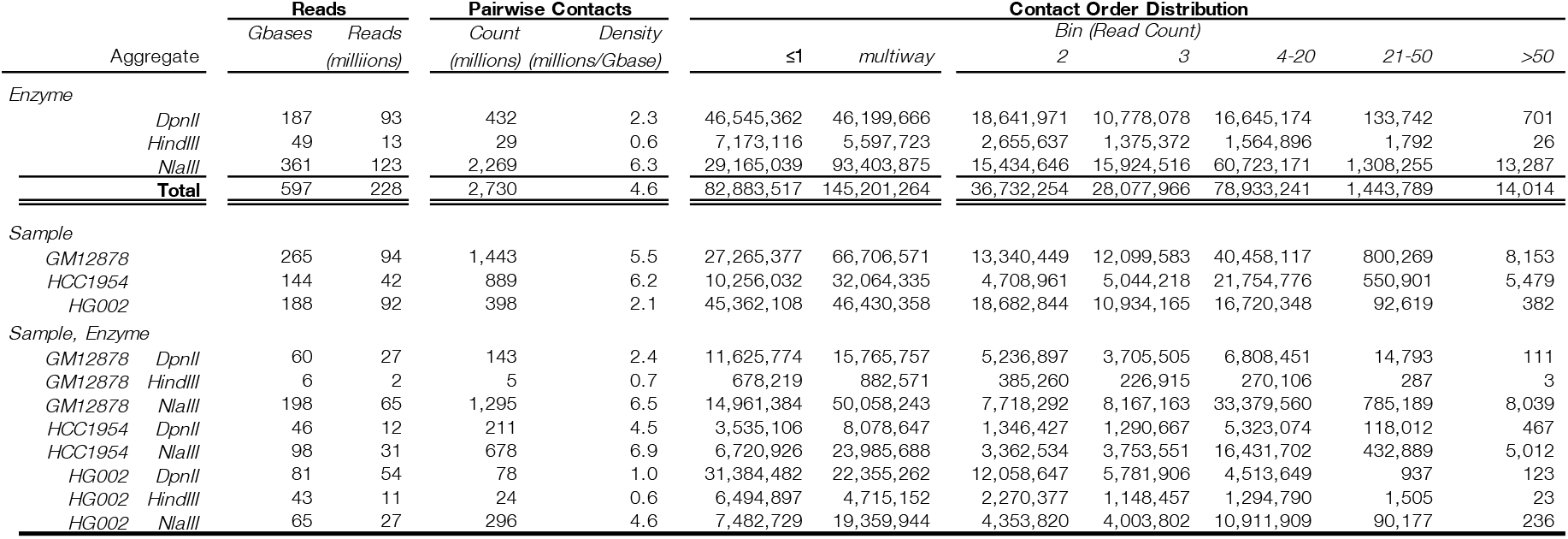
Pore-C sequencing runs in aggregate. Summary data aggregated across all runs by either sample, or by sample and enzyme.

**Table S2.**
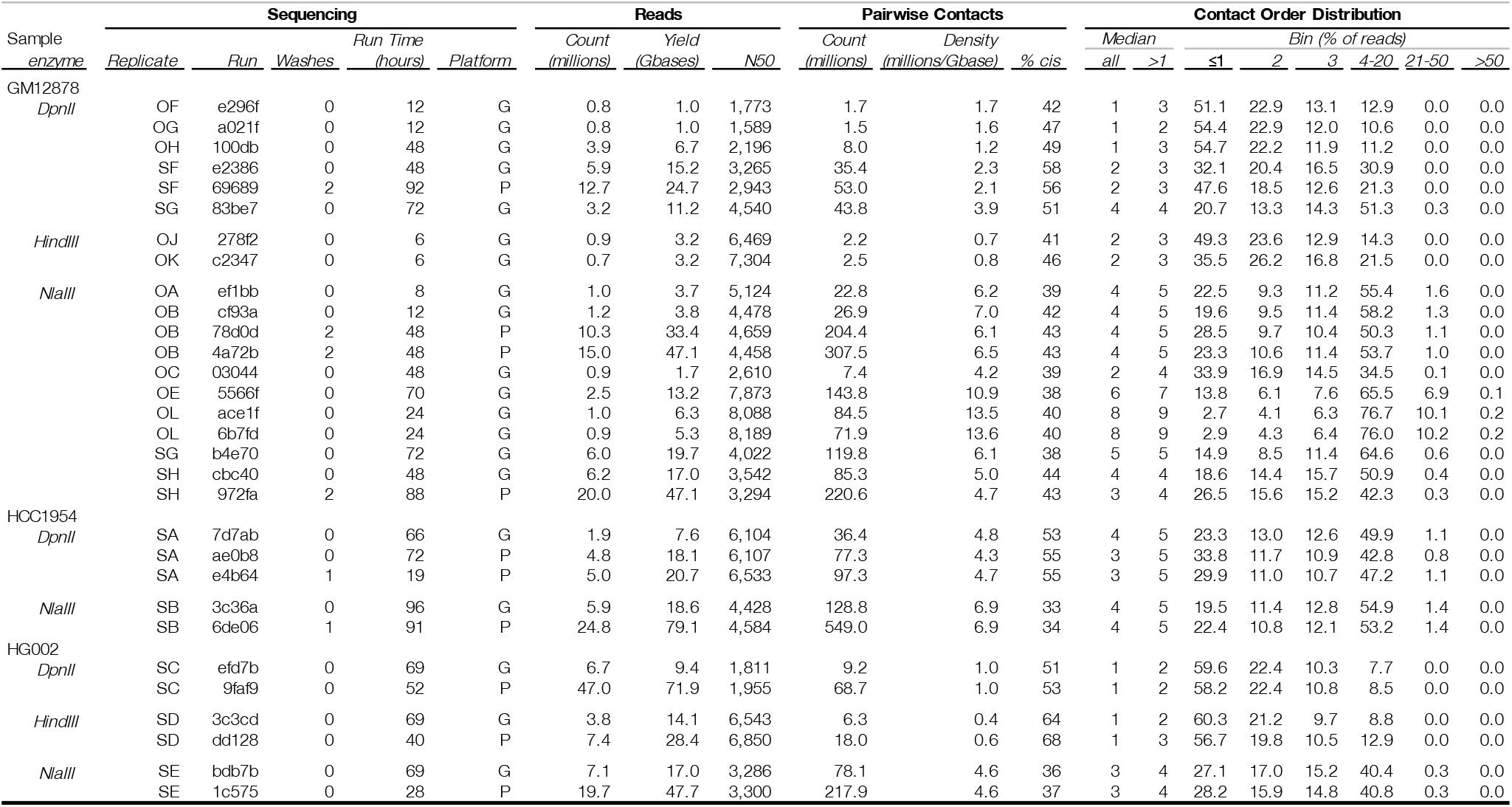
Pore-C sequencing runs. Per-run statatistics and results for all Pore-C sequencing runs used in this publication.

**Table S3.** Commands and parameters used in this study. A table of all the computational tools used in this study to process Pore-C data. These parameters are the ones implemented in the snakemake workflow.

| tool | ver | link | parameters | purpose |
| --- | --- | --- | --- | --- |
| pore_c refgenome virtual-digest | 0.1 | https://github.com/nanoporetech/pore-c/ | default parameters | generates an in-silico restriction map for assignment of alignments to restriction fragments. |
| pore_c reads catalog | 0.1 | https://github.com/nanoporetech/pore-c/ | default parameters |  |
| pore_c alignments parse | 0.1 | https://github.com/nanoporetech/pore-c/ | default parameters | conducts alignment filtering from the raw bam file using the fragDAG algorithm. |
| pore_c to-matrix | 0.1 | https://github.com/nanoporetech/pore-c/ | default parameters | Computes pairwise contacts from multiway contacts in the filtered alignment table data and writes pairwise contacts to the standard pairs file format. |
| bwa | 0.7.17 | https://github.com/lh3/bwa/ | `bwasw -b 5 -q 2 -r 1 -T 15 -z 10` | alignment of pore-c monomers to a genome |
| cooler load | 0.8.6 | https://github.com/mirnylab/cooler/ | `--input-copy-status unique -f coo` | creates a 1kb binned cooler file from a pairs file created from a decomposed pore-C multiway contact data set |
| cooltools diamond-arrowhead | 0.2.0 | https://github.com/mirnylab/cooltools/ | `3 5 10 15 --window-pixels --append-raw-scores` | calculates the insulation score vector used to identify topologically associated domains. |
| cooltools call-compartments | 0.2.0 | https://github.com/mirnylab/cooltools/ | default parameters | calculates the balanced matrix eigenvectors which indicate the A and B compartments |
| cooler merge | 0.8.6 | https://github.com/mirnylab/cooler/ | default parameters |  |
| cooler balance | 0.8.6 | https://github.com/mirnylab/cooler/ | `--blacklist blacklist.bed --convergence-policy store_nan --max-iters 10000` | balances the pairwise contact matrix |
| cooler zoomify | 0.8.6 | https://github.com/mirnylab/cooler/ | default parameters | generates a balanced multi-resolution cooler from a 1kb binned matrix file. |
| wtdbg2 | 2.5 | https://github.com/ruanjue/wtdbg2/ | `--edge-min 4 --force --kmer-psize 19 --kmer-subsampling 4 --preset ont --rescue-low-cov-edges --tidy-reads 5000` | generates a genome assembly from ONT long read sequencing data |
| racon | 1.4.7 | https://github.com/lbcb-sci/racon/ | `--error-threshold 0.3 --gap -8 --include-unpolished --match 5 --num-corrections 3 --quality-threshold -1 --window-length 1000` | assembly polishing |
| Medaka | 0.9.2 | https://github.com/nanoporetech/medaka/ | `-b 100 -m r941_min_high` | assembly polishing |
| SALSA2 | 2.2 | https://github.com/marbl/SALSA/ | `--iter 10 --clean yes -p yes` | uses pore-C pairwise data to fix and scaffold the wtdbg2 assembly |

**Fig. S2.**
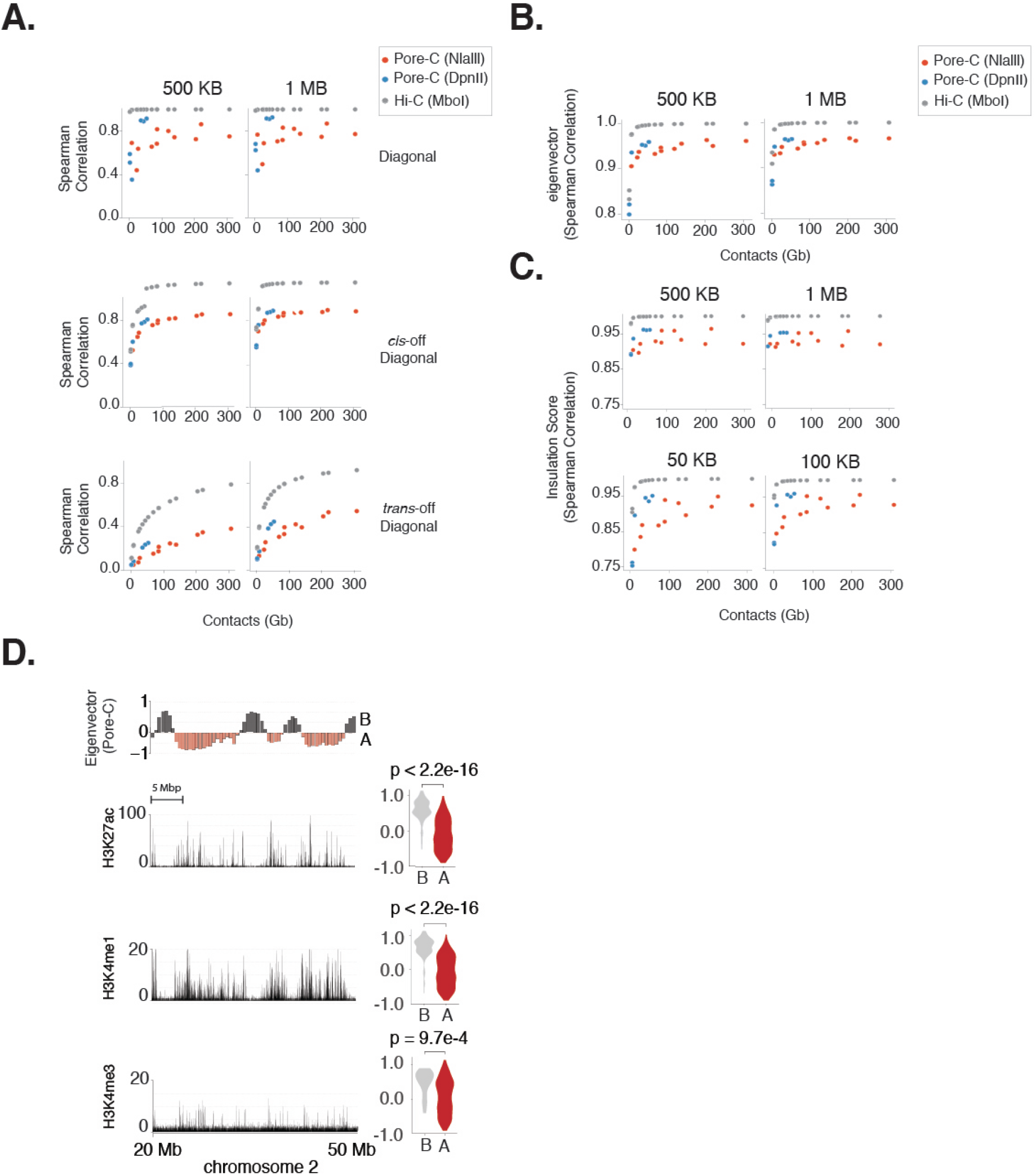
Genomic Compartmentalization. Similarity to the gold standard Hi-C dataset to individual Pore-C sequencing runs. Grey dots represent the upper bound of similarity of a Pore-C run of the indicated size, and is produced by downsampling the gold standard data to the same size as the indicated Pore-C run, and comparing it to the whole Hi-C dataset. Similarity is assessed using (A) Whole unbalanced matrix correlation along the diagonal (top), between all *cis* contacts not on the diagonal (middle), and all *trans* contacts (bottom), (B) using correlation of the compartmental eigenvector values of individual runs to the gold standard Hi-C data set, or (C) Correlation of the insulation scores of the individual runs with the gold standard Hi-C data set. (D) Correlation of the eigenvector compartment score with other indicators of active and inactive chromatin state along a 30 Mbp window in chromosome 2 binned at 500 Kbp resolution, including ChIP-seq peaks for the histone marks H3K27ac, H3K4me3 and H3K4me1. At right, violin plots indicate the correlation of the A and B compartments with the presence of ChIP-seq peaks.

## Word Counts

This section is *not* included in the word count.

### Notes on JPhysD article

- Abstract: 150 words, unreferenced
- Main text: short format 4500 words, long is 8500 words.
- No limit in number of figures

## Statistics on word count

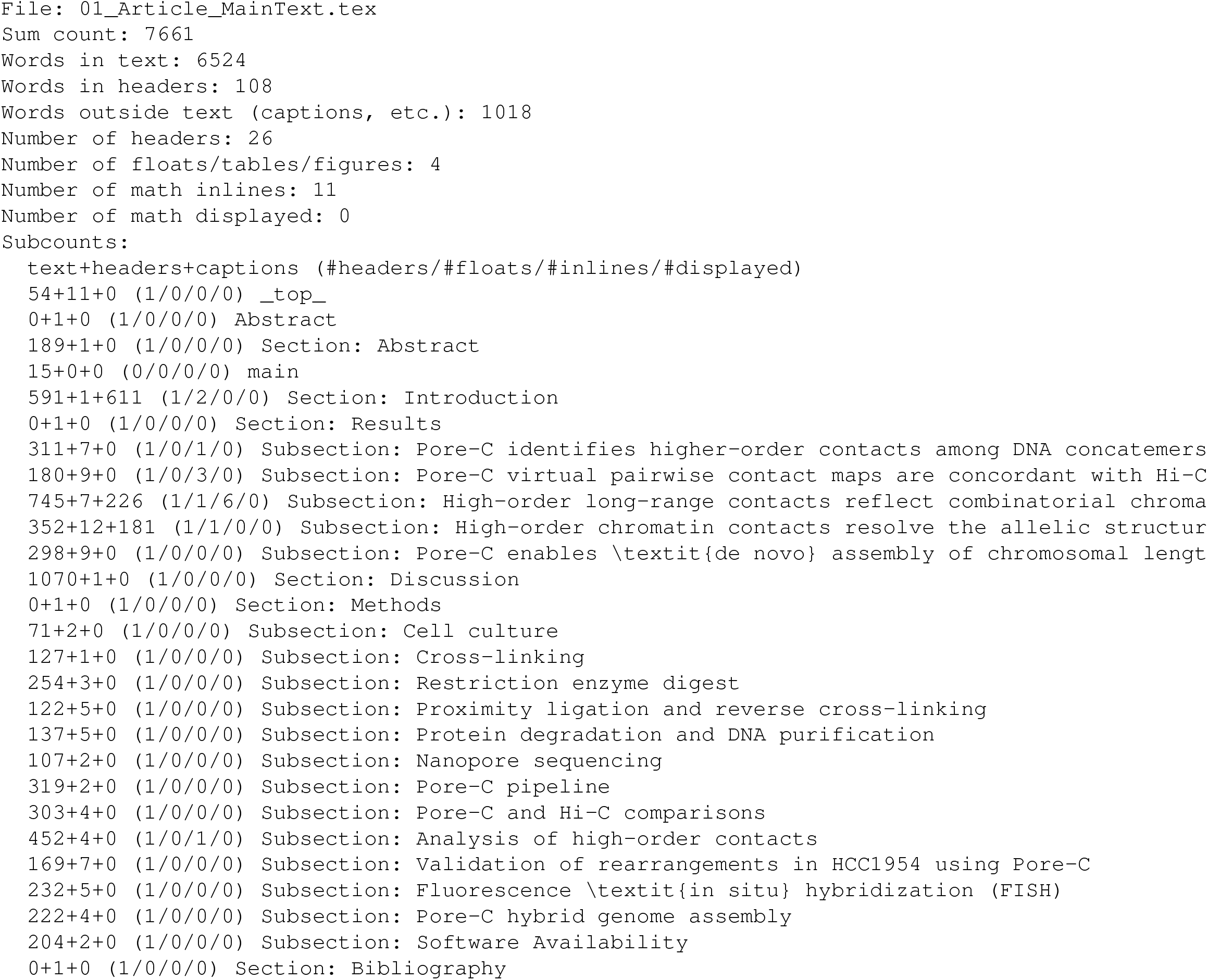

